# A simple method for gene expression in endo- and ectodermal cells in mouse embryos before neural tube closure

**DOI:** 10.1101/2020.05.14.086330

**Authors:** Yusuke Kishi, Yurie Maeda, Naohiro Kuwayama, Yukiko Gotoh

## Abstract

The lack of a widely accessible method for expressing genes of interest in wild-type embryos is the fundamental obstacle in understanding genetic regulation during embryonic development. In particular, only a few methods are available for introducing gene expression vectors into cells prior to neural tube closure, a period for the drastic development of many tissues. In this study, we present a simple technique for transferring vectors into the endo- and ectodermal cells of mouse embryos at E7.0 or E8.0 via in utero injection, without any specialized equipment. Using this technique, introduction of retroviruses can facilitate the labeling of cells in various tissues, including the brain, spinal cord, epidermis, and digestive and respiratory organs. As such, this technique can aid in analyzing the roles of genes of interest during endo- and ectodermal development prior to neural tube closure.

## Introduction

To understand the molecular mechanisms underlying mammalian development, it is necessary to manipulate the expression of the genes of interest in mouse embryos. The most conventional method for such manipulation involves generating gene-targeted knock-in or knock-out mice and crossing them with specific Cre- or CreERT2-expressing mouse strains ^1–3^. Although the advent of genome editing tools, such as the transcription activator-like effector nuclease and clustered regularly interspaced palindromic repeat (CRISPR)/CRISPR-associated protein 9 systems ^4–7^, has made it easier to develop transgenic mice, these tools are expensive, labor-intensive, and inaccessible to many biologists owing to the need for specialized equipment, such as micromanipulators or electroporators.

An alternative approach for gene manipulation is the direct delivery of vectors into wild-type embryos ^8^. In the central nervous system, in utero injection of a DNA plasmid, followed by electroporation or introduction of a viral vector into the ventricle is a widely utilized method for embryos after embryonic day 10.5 (E10.5) ^9–11^. In utero injections of viral vectors were also applied to the epithelium and nervous system after E9.5 ^12–15^. For analyzing the role of genes of interest at E7 or E8, a period of rapid organ development, ultrasound-guided method is useful to introduce viral vectors into embryos. Although previous methods for E8.5 embryos resulted in low survival rates and abnormalities in many surviving embryos^16^, a recent method for E7.5 embryos, called NEPTUNE, greatly improved the survival rate of embryos ^17^. As the neural tube is still open at E7.5, a viral vector injected into the amniotic fluid can access the neuroepithelial cells in addition to the epithelial and endodermal tissues^17^. The high efficiency of gene transduction achieved by the NEPTUNE-mediated knockdown of *Olig2* or *Sptbn2* resulted in phenotypes similar to those of their knockout embryo counterparts ^17^. However, this approach requires specialized and expensive equipment as well as an ultrasound microscopy system.

In this study, we developed a method to introduce retroviruses into the amniotic fluid at E7.0 and E8.0 for gene manipulation during organ morphogenesis without an ultrasound microscopy system. Most embryos survived and exhibited normal development. We confirmed ectopic gene expression in the endodermal and ectodermal tissues, similar to that observed with the NEPTUNE system. In addition, we demonstrated the possibility of introducing plasmids via electroporation, although this requires further investigation. Therefore, our method is useful for analyzing the functions of genes of interest and beneficial to the broader scientific community.

## Results

### Development of a simple and efficient method to introduce a retrovirus in mouse embryos prior to neural tube closure

We hypothesized that it would be feasible to introduce a reagent into amniotic fluid without using an ultrasound microscopy system by regulating the position of the capillary tip in the uterus. To this end, we employed optical fibers to visualize the inner structure of the uterus, including the amnion and embryo (Figure 1a–c). Optical fiber-based visualization of the ventricles of embryos from E9.5, has been previously described for targeted DNA injection and electroporation following neural tube closure (Figure 1c)^18^. We found that lighting with this system was adequate to enable the injection of the dye inside the amnion when positioning the tip of the capillary at the center of the uterus (see Methods), although it was not possible to directly visualize the amnion within the uterus. To precisely control the developmental stage of embryos, mice were mated for only 2 h, and females with a confirmed vaginal plug were used for in utero injection. We first injected carboxyfluorescein succinimidyl ester (CFSE), a cell-permeable dye that becomes fluorescent following its reaction with intracellular proteins^19,20^, into the uterus at E8.0. To minimize the potential damage caused by the injection, we inserted a sharpened capillary at the center of the uterus at an angle perpendicular to the uterine wall while it was positioned (Figure 1d). We injected a small volume (approximately 0.5 μL) of CFSE solution at a depth of 1–2 mm below the uterine wall, which resulted in reproducible labeling of the embryos. The duration of surgery is relatively short (approximately 20 min) because of the simplicity of the method, which may contribute to minimal damage to the embryos. After three days (at E11.0), fluorescence microscopy without sectioning revealed that 77% of the injected embryos were labeled with CFSE (Figure 1e–f). Approximately 80% of the embryos survived without gross developmental malformations at E11.0. We also injected CFSE into the amniotic fluid at E7.0, which yielded a high rate of embryo labeling (64%) and a high rate of survival without gross developmental malformations (89%) at E9.0 (Figure 1e–f).

**Figure 1.**
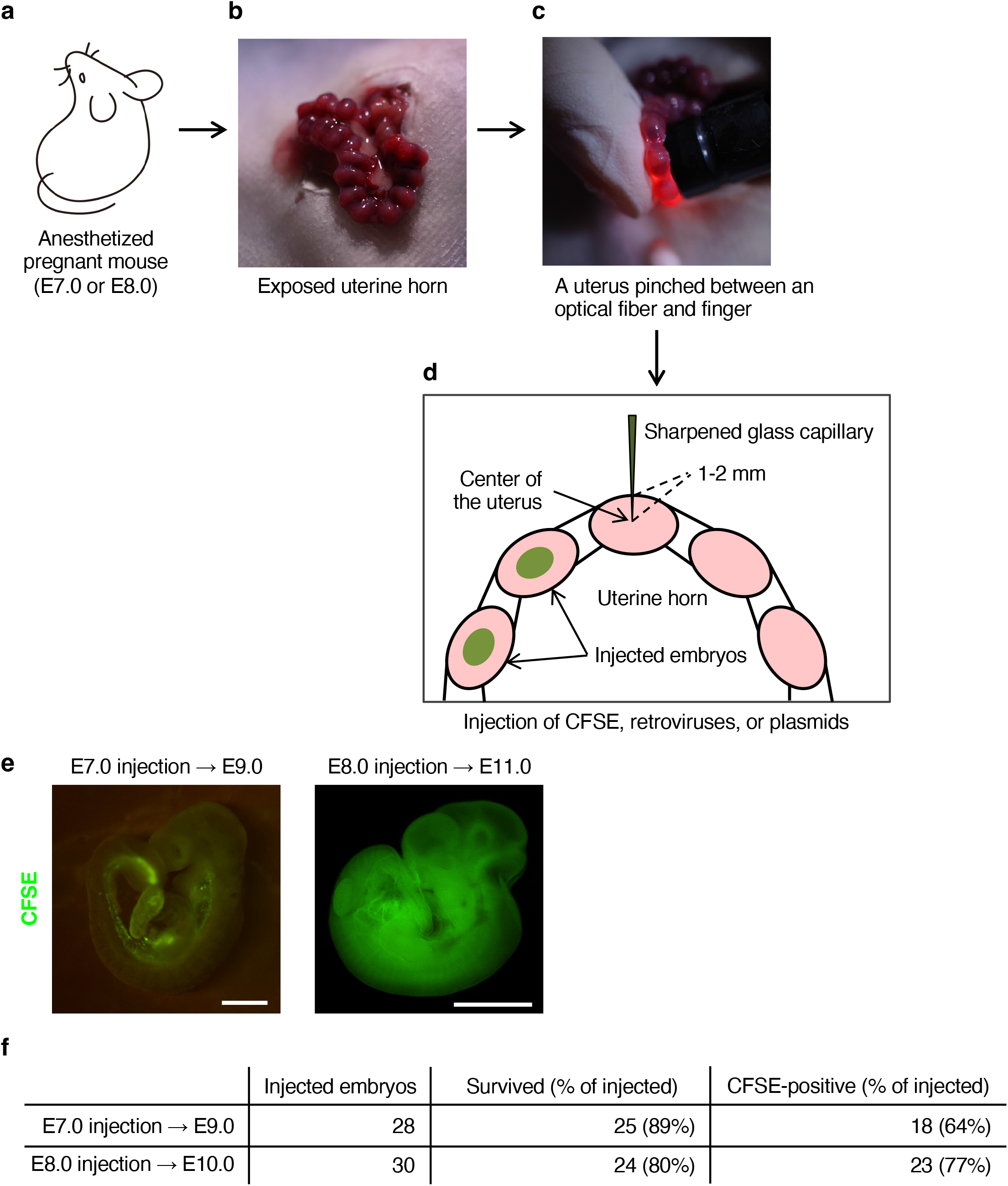
In utero injection into mouse embryos before neural tube closure. (**a–b**) Exposure of the uterine horn of an anesthetized pregnant mouse at embryonic day 7 (E7.0) or E8.0. (**c–d**) Pinching of the uterus between an optical fiber and finger (**c**) for injection via a sharpened glass capillary of a solution colored with a dye (FastGreen), containing either carboxyfluorescein succinimidyl ester (CFSE), a retrovirus, or a plasmid (**d**). (**e**) Embryos at E7.0 (left) or E8.0 (right) were injected with CFSE, dissected out from the uterus at E9.0 or E11.0, respectively, and fixed for the visualization of CFSE fluorescence with a fluorescence stereo microscope. Scale bars, 0.5 mm (left) or 1 mm (right). (**f**) Survival and labeling rates for the in utero introduction of CFSE into the embryos of three pregnant mice in each condition as described in (**e**).

Subsequently, we used retroviral vectors for gene introduction using this method. We initially injected a concentrated suspension of a retrovirus encoding green fluorescent protein (GFP) under the control of the phosphoglycerate kinase (*PGK*) promoter, which is the same promoter utilized in the NEPTUNE system, into the uterus at E8.0. Our results revealed that 89% of embryos survived without gross abnormalities, and 74% of embryos expressed GFP at E11.0 (Figure 2a and c). Injection of the retrovirus at E7.0, also induced GFP expression at E11.0, although the labeling rate was lower (Figure 2b–c). Flow cytometric analysis of the cortical tissue obtained from E12.0 embryos that had been injected with the retrovirus encoding GFP at E8.0 showed that approximately 70% of the cortical cells were positive for GFP in successful cases (Figure 2d). Immunohistochemical analysis of sagittal sections at E10.0 embryos that had been injected with the retrovirus encoding GFP at E8.0 revealed that most tissues exposed to amniotic fluid expressed GFP (Figure 2e–g). The labeled tissues included the central nervous system (forebrain, midbrain, hindbrain, and spinal cord) as well as the epidermis and endodermal tissues, such as digestive and respiratory organs. This suggests that our method is suitable for gene introduction into ectodermal and endodermal tissues at the early stages of mouse development.

**Figure 2.**
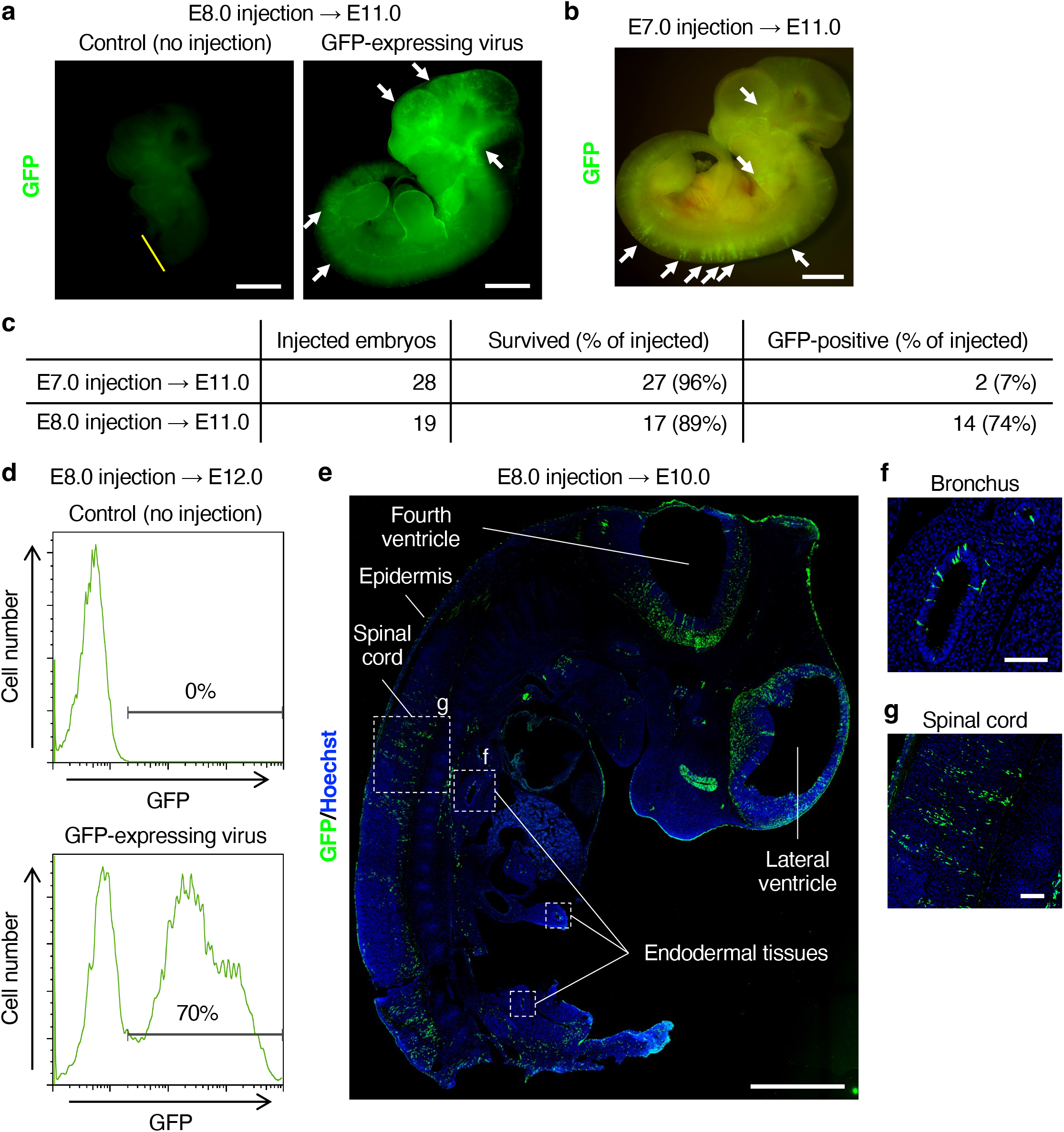
In utero gene introduction via retroviral infection in embryos before neural tube closure. (**a**) Embryos within the same uterus were administered with a retrovirus encoding green fluorescent protein (GFP) under the control of the phosphoglycerate kinase (*PGK*) promoter at E8.0. The embryos were subsequently dissected from the uterus at E11.0, fixed, and examined for GFP fluorescence using a fluorescence stereo microscope. The posterior half of the control embryo was excised at the yellow line for identification purposes. Arrows indicate the GFP-expressing regions. Scale bar, 1 mm. (**b**) An embryo at E7.0 was administered with a retrovirus as in (**a**) and examined for GFP fluorescence at E11.0. Arrows indicate the GFP-expressing regions. Scale bar, 1 mm. (**c**) Survival and labeling rates for embryos as in (**a**) and (**b**). (**d**) Flow cytometric analysis of the gene introduction efficiency. Neocortical cells dissociated from E12.0 embryos administered the retrovirus for GFP at E8.0 were subjected to flow cytometric analysis of GFP expression. This embryo exhibited the highest rate of GFP-positive cells in our experiments. (**e**) A sagittal section of an E10.0 embryo injected at E8.0 as in (**a**) was subjected to immunohistofluorescence staining with anti-GFP antibodies. Nuclei were counterstained with Hoechst 33342. Representative infected tissues are indicated. Scale bar, 1 mm. (**f**–**g**) High magnification images of the bronchus (**f**) and spinal cord (**g**) as shown in (**e**). Scale bar, 100 μm.

### Possible application of plasmid introduction via electroporation

Given that in utero electroporation of plasmids is generally more efficient and results in higher expression levels than retroviral infection as a method of gene introduction, we next employed our new method to inject a plasmid encoding mCherry and performed in utero electroporation using electrodes (disc size, 2.5 by 4 mm) positioned on each side of the uterus at E7.0 or E8.0. The proportions of mCherry-expressing embryos at E9.0 and E10.0 were 19% and 40% among those electroporated at E7.0, and E8.0, respectively (Figure 3a and b). The expression pattern of mCherry varied among embryos, likely because of differences in the orientation of the electrode relative to the embryo during electroporation. Most embryos expressed mCherry in the somites (Figure 3a), whereas only one of them expressed mCherry in the midbrain and hindbrain (Figure 3c). To date, no embryos have expressed mCherry in the forebrain. These results demonstrate that it is possible to perform gene introduction by in utero electroporation in mouse embryos at E7.0 or E8.0 using our method, although further improvements are necessary to allow the targeting of a specific tissue.

**Figure 3.**
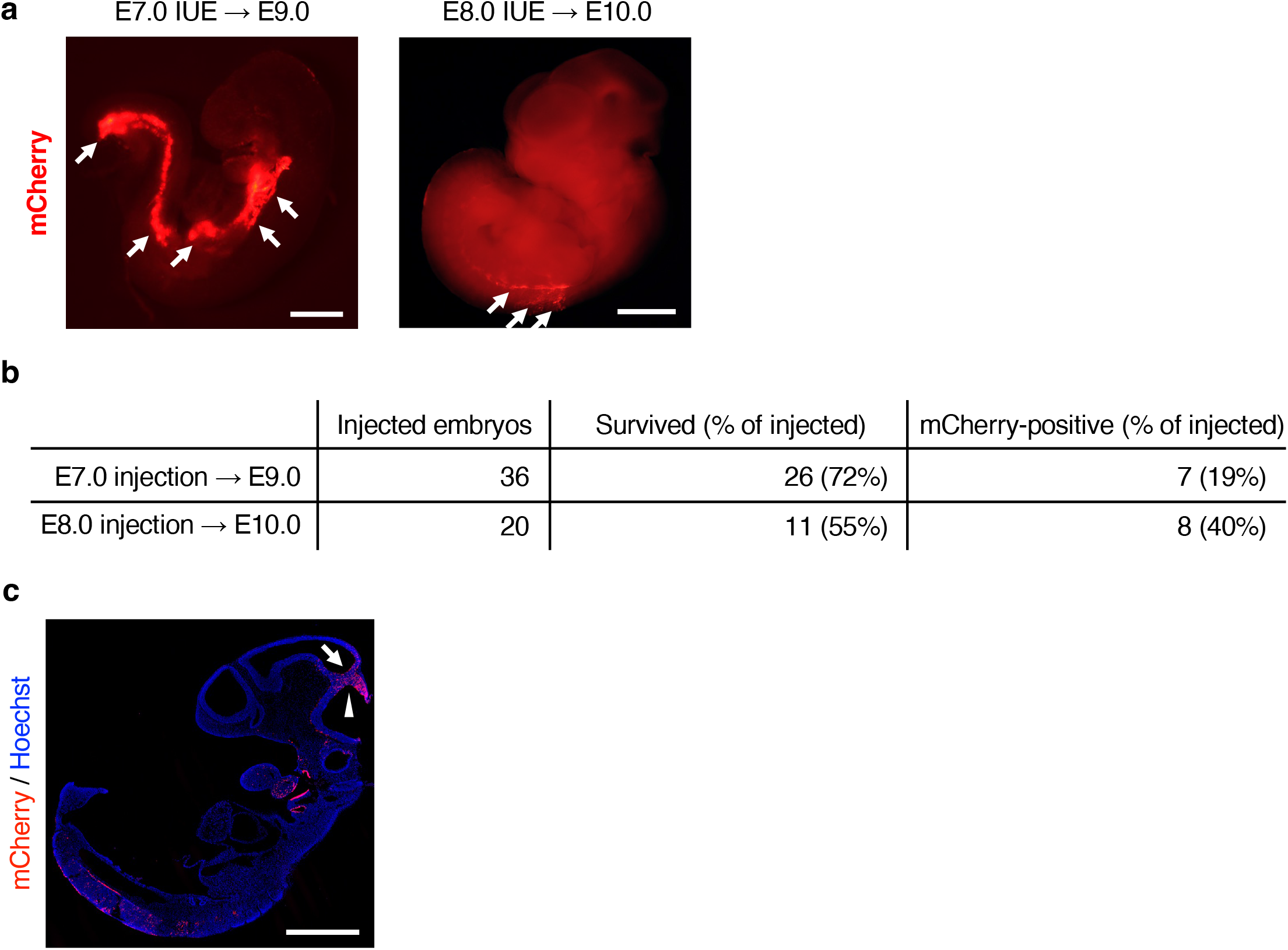
In utero electroporation into mouse embryos before neural tube closure. (**a**) Embryos at E7.0 (left) or E8.0 (right) were injected with a plasmid encoding mCherry and subjected to in utero electroporation. They were subsequently harvested from the uterus at E9.0 or E10.0, respectively, fixed, and examined for mCherry fluorescence using a fluorescence stereo microscope. Arrows indicate the mCherry-expressing regions. Scale bars, 0.5 mm (left) or 1 mm (right). (**b**) Survival and labeling rates for the in utero electroporation of embryos as described in (**a**). (**c**) A sagittal section of an E9.0 embryo electroporated at E8.0 as in (**a**) was subjected to immunohistofluorescence staining with anti-mCherry antibodies. Nuclei were counterstained with Hoechst 33342. Arrow and arrowhead indicate the mCherry-expressing regions in third and fourth ventricles, respectively. Scale bar, 1 mm.

## Discussion

Manipulation of gene expression is necessary for the detailed investigation of molecular mechanisms underlying embryonic development. However, in the central nervous system, existing methods, such as the injection of viruses or plasmids for in utero electroporation into the ventricles, can only be applied after neural tube closure ^9–11^. The gene introduction method developed in the present study has several advantages over other methods that can be used before neural tube closure, such as whole-embryo genomic manipulation (gene targeting or transgene introduction) and ultrasound-guided injection of viruses into the mouse nervous system. Genomic manipulation allows the precise expression of a gene of interest in a temporally and spatially regulated manner; however, it requires a specific Cre-expressing mouse strain or knowledge of specific distal elements to control expression, and a significant investment of time, cost, and space to establish and maintain a transgenic mouse strain. The NEPTUNE method, which uses the ultrasound-guided injection of viruses, addresses many weaknesses of genomic manipulation and allows for the analysis of a gene of interest in less time, space, and animals ^17^. However, it still requires specific equipment, such as ultrasound microscopy.

Our present method, which is based on the same principle as the NEPTUNE method, introduces a chemical and virus into embryos by injection into the amniotic fluid before neural tube closure; however, it is possible to introduce it without ultrasound microscopy. We also propose the possibility of introducing plasmids by electroporation, although further improvements, such as electroporation at multiple electrode orientations to a single embryo, are necessary. The gene transfection efficiency of the retrovirus using our method was lower than that using the NEPTUNE method (approximately 70% in our best case and >99% in NEPTUNE) ^17^. This discrepancy may be attributed to the relatively imprecise targeting of the injection site in our method, which did not utilize ultrasound guidance as well as differences in the types and preparation methods of the viruses used. While Mangold et al. reported that an injection of more than 0.25 μL of lentivirus solution decreased the survival rate, our method allows for the injection of approximately 0.5 μL of retrovirus solution^17^. The simplified procedure of our method, with a shorter surgery time (approximately 20 min) and no requirement for pregnancy and stage verification due to the limited mating time, may contribute to a higher survival rate of embryos. However, transfected tissues were comparable between the two methods. As such, our method is a useful tool for testing and screening various genes for their specific effects on development and will be beneficial to the broader scientific community.

## Materials and Methods

### Mouse maintenance

All animal studies were conducted in accordance with the protocols approved by the Animal Care and Use Committee of the University of Tokyo. Mice were housed in a temperature- and humidity-controlled environment (23 ± 3 °C and 50 ± 15%, respectively) under a 12-h light/dark cycle. Animals were housed in sterile cages (Innocage, Innovive) containing bedding chips (PALSOFT, Oriental Yeast), with two–six mice per cage, and provided irradiated food (CE-2, CLEA Japan) and filtered water ad libitum.

### Plasmid construction

cDNA encoding mCherry was amplified via polymerase chain reaction, and its coding sequence was confirmed via DNA sequencing. The cDNA was then subcloned into the pCAGEN vector, kindly provided by C. L. Cepko, to generate the pCAGEN-mCherry plasmid.

### Retrovirus production and concentration

PLAT-GP cells, a gift from T. Kitamura, were harvested on a 10-cm diameter dish (55 cm^2^ surface area) at 80–90% confluency in Dulbecco’s Modified Eagle Medium (Sigma) containing 10% fetal bovine serum and 1% penicillin–streptomycin (Thermo Fisher Scientific). The cells were transfected with polyethyleneimine (PEI; Polysciences). A mixture of 12 μg of the validated retroviral vectors pSIREN_EGFP-siLuc (Addgene #122287) ^21^ expressing GFP under the control of *PGK* promoter and 9 μg of a VSVG plasmid, kindly provided by H. Song, were mixed in a 1.5-mL tube with 500 μL of OPTI-MEM (Thermo Fisher Scientific). Then, 63 μL of 1 mg/mL PEI in water in another 1.5-mL tube with 50 μL of OPTI-MEM was shaken vigorously and incubated at room temperature (20–25 °C) for 5 min. The plasmid and PEI solution were mixed, shaken vigorously, and incubated at room temperature for 15 min. The mixture was then added to the PLAT-GP cells.

The medium containing the transfection reagents was replaced with 10 mL of fresh medium without penicillin–streptomycin, and the conditioned medium collected 72 h after the onset of transfection was cleared of debris via centrifugation at 2,500 × *g* for 5 min, passed through a 0.45-μm filter, and centrifuged again at 250,000 × *g* for 90 min at 4 °C. After roughly removing the supernatant via decantation, the pellet was resuspended with the remaining medium, transferred to a new 0.6-mL tube, and centrifuged at 20,000 × *g* for more than 12 h at 4 °C. The supernatant was completely removed via pipetting, and the pellet was resuspended in 10 μL of phosphate-buffered saline. The 1,000-fold concentrated retrovirus was stained with 0.01% FastGreen (Chroma Gesellschaft), rapidly frozen with liquid nitrogen, and stored at –80 °C. The viral titer was not determined.

### Preparation of pregnant mice and capillary filled with virus, CFSE, or plasmid

JCL:ICR (CLEA Japan) mice were used for in utero injection experiments. To ensure precise control of the embryonic stage, mating was limited to a 2-h period, and the presence of a vaginal plug was used to confirm the onset of gestation (E0.0).

Glass capillaries (GD-1; Narishige) were pulled using a PC-100 (Narishige) puller, roughly cut with forceps, and shaped with an EG-45 (Narishige) grinder to achieve a tip diameter of 70–80 μm and an angle of approximately 30°. These capillaries were then backfilled with thawed viral, CFSE, or plasmid solution using a microloader (Eppendorf) and attached to a FemtoJet (Eppendorf) injector. Any remaining air at the tip was eliminated via flushing.

### In utero injection and electroporation before neural tube closure

Pregnant mice were administered a sodium pentobarbital-based anesthetic, and the uterine horn was exposed and pinched between a flexible fiber-optic cable and a finger ^18^ (Figure 1c). Lighting from the fiber-optic cable helped in the visualization of the internal uterine structure, but did not enable the identification of the embryo from the amnion. CFSE (1 mM; Sigma), plasmid DNA (1 μg/μL), or retroviruses, each mixed with FastGreen, were injected at an angle perpendicular to the center of the uterus using a FemtoJet (Eppendorf) at 100–150 hPa pressure. An injection depth of 1–2 mm below the uterine wall permitted consistent introduction of capillary contents into the amniotic fluid. After visually confirming the presence of FastGreen at the center of the uterus, the injection was halted, and the capillary was immediately removed. Generally, approximately 0.5 μL of the reagent was injected, although this amount varies based on the embryo. For electroporation, the uterus was held between tweezer-type electrodes (CUY652P2.5×4; Nepa Gene) and electroporated with a NEPA21 system (Nepa Gene), according to the following program: three poring pulses of 30 V with a 15-ms pulse width and 450-ms pulse interval and 10% decay (+ pulse orientation), and two transfer pulses of 10 V with a 50-ms pulse width and 450-ms pulse interval and 10% decay (± pulse orientation). After injection and electroporation, the uterine horn was returned to its original location. Pregnant mice were allowed to recover on a heating plate maintained at 38 °C until they regained consciousness following anesthesia. Following the designated period of observation, the embryos were removed and examined using a fluorescence stereo microscope equipped with a 2.0 × objective lens manufactured by Nikon (SMZ18).

### Flow cytometry

Flow cytometry was performed using a previously established protocol ^22^. Specifically, the neocortices of the ICR embryos were dissected and subjected to enzymatic digestion using Neuron Dissociation Solution (FUJIFILM Wako). To quantify the efficiency of gene introduction, cells were analyzed using the FACSAria instrument (Becton Dickinson) via direct flow cytometry.

### Immunohistofluorescence analysis

Immunofluorescence staining was performed as described previously ^23^. A 1:2,000 dilution of the anti-GFP antibody (Abcam; ab13970) and a dilution 1:000 of the anti-DsRed antibody (Takara; 632496) for mCherry were used, and the nuclei were counterstained with a 1:10,000 dilution of Hoechst 33342. Images were acquired using a laser-scanning confocal microscope (TSC-SP5; Leica) and analyzed using the ImageJ software (NIH).

## Acknowledgments

We would like to thank T. Shimogori (RIKEN) as well as M. Miura, Y. Yamaguchi, K. Hashimoto, and H. Miyazawa (The University of Tokyo) for technical advice on in utero injection before neural tube closure; H. Song (Johns Hopkins University School of Medicine, Baltimore, MD) for providing the plasmid encoding VSVG; T. Kitamura (The University of Tokyo) for providing PLAT-GP cells; C. L. Cepko (Harvard Medical School) for providing the pCAGEN plasmid; The One-Stop Sharing Facility Center for Future Drug Discoveries (The University of Tokyo) for providing FACS; R. Nagayoshi, and Y. Kakeya (The University of Tokyo) for technical assistance; and members of Gotoh and Kishi laboratories for helpful discussion. This research was supported by AMED-CREST (JP22gm1310004 to Y.G.), AMED-PRIME (JP22gm6110021 to Y.K.), MEXT/JSPS KAKENHI (JP22H00431 to Y.G.; JP20H03179, JP21H00242, and JP22H04687 to Y.K.), Takeda Science Foundation, and the SECOM Science and Technology Foundation.

## Author Contributions

Y.K. designed and supervised the study and wrote the manuscript. Y.G. supervised the study. Y.K., Y.M., and N.K. performed the experiments and analyzed the data.

